# Decoding region-level visual functions from invasive EEG data

**DOI:** 10.1101/2024.04.02.587853

**Authors:** Xin-Ya Zhang, Hang Lin, Zeyu Deng, Markus Siegel, Earl K. Miller, Gang Yan

## Abstract

Decoding vision is an ambitious task as it aims to transform scalar brain activity into dynamic images with refined shapes, colors and movements. In familiar environments, the brain may trigger activity that resembles specific pattern, thereby facilitating decoding. Can an artificial neural network (ANN) decipher such latent patterns? Here, we explore this question using invasive electroencephalography data from monkeys. By decoding multiregion brain activity, ANN effectively captures individual regions’ functional roles as a consequence of minimizing visual errors. For example, ANN recognizes that regions V4 and LIP are involved in visual color and shape processing while MT predominantly handles visual motion, aligning with regional visual functions evident in the brain. ANN likely reconstructs vision by seizing hidden spike patterns, representing stimuli distinctly in a two-dimensional plane. Furthermore, during the encoding process of transforming visual stimuli into neuronal activity, optimal performance is achieved in regions closely associated with vision processing.

## Introduction

In recent years, there has been a growing interest in the fields of mind reading and encoding, spurred by the rise of brain-inspired machines [1, 2, 3, 4, 5]. Decoding visual information, in particular, poses a formidable challenge, as it involves converting the scale of neuronal spikes into a high-dimensional image featuring color schemes, brightness and shape [6, 7]. This challenge is analogous to attempting to represent a broad range of possibilities with a limited set of options. Metaphorically speaking, it’s akin to trying to capture the full spectrum of colors using just a handful of crayons.

The potential to reconstruct visual information from the brain may rely on discerning specific neuronal spike patterns in familiar environments, facilitating the transformation of spatio-temporal features into meaningful images. While diverse architectures of artificial neural network (ANN)-based machines have been proposed, interpretability plays a crucial role in determining their adequacy as brain models [8, 9, 10, 11, 12, 13]. Hence, it is important not only to improve the accuracy of visual decoding but also to reveal the underlying mechanism in the mapping from visual stimuli to brain activity. Does the ANN-based machine learn like a brain does? Previous studies have found such convergence to some extent in task-oriented attention [14, 15], such as implicit attention in CNNs [16] and LSTMs [17]. However, compelling evidence is still lacking regarding the functional convergence between the brain and machine, especially through the error-minimization strategy widely used in ANNs. This gap extends to the consistency of encoding and decoding models effectively mirroring region-specific or neuron-cluster functions, particularly in the domain of brain-computer interfaces.

The central hypothesis guiding our study posits that neurons in specific brain regions would collectively exhibit similar spike patterns when the brain operates in a familiar environment. To explore this, we analyzed an ideal invasive electroencephalography (iEEG) dataset [18], which involves two rhesus monkeys trained on a categorization task over years and offers superior spatial and temporal resolution compared to fMRI data [19, 20, 21]. During the experiments, when trained monkeys saw a series of stimulus videos, trial-by-trial neuronal activity was recorded across six brain regions: frontal cortex (lateral prefrontal cortex: PFC and frontal eye fields: FEF), parietal cortex (lateral intraparietal cortex: LIP), and occipitotemporal cortex (inferotemporal cortex: IT, visual area: V4, and middle temporal cortex: MT).

The spike recording collected from multiple brain regions offers a valuable opportunity to assess a machine’s ability to comprehend intricate neural coding and capture regional functions in response to efforts aimed at minimizing visual errors. Considering that a specific function may be confined to particular areas, it is crucial to assess the feasibility of visual decoding from limited regions. In the experiment data, regions LIP, IT, and V4 participate in visual processing, and MT is involved in visual motion. Additionally, cognitive regions PFC and FEF contribute to decision-making processes and may influence the integration of visual information.

The stimulus video presented to two rhesus monkeys comprised three main phases: a 0.5-second fixation period, a 1-second cue display, and a 3-second stimulus presentation, as depicted in Fig. 1A. During each trial, the cue image was selected from four possible shapes, while the continuous stimulus consisted of combinations of four possible colors and four possible directions, resulting in a set of 16 types of stimuli (Fig. 1B). Consequently, there were 64 types of stimulus video combinations in total. The initial position and speed of stimulus dots varied from trial to trial. The rhesus monkeys executed a saccade based on the task cue, indicating whether color or direction of movement was relevant. Notably, our study did not involve the categorization task; instead, our focus was directed towards examining the relationship between neuronal activity and the stimulus video.

**Fig. 1.**
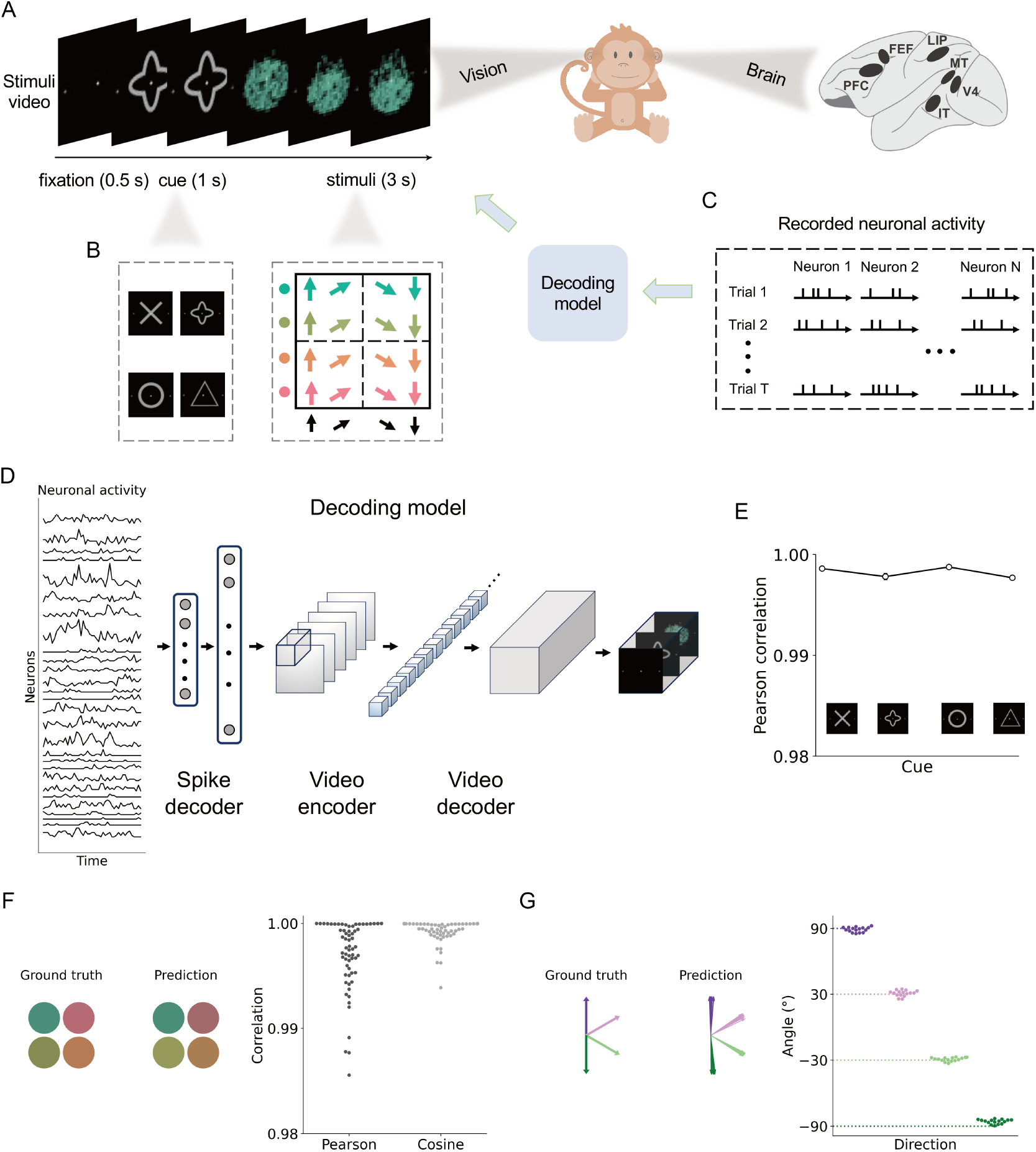
Decoding dynamic visual stimuli from monkey iEEG data. (**A**) Neuronal activity was recorded trial-by-trial in six brain regions when monkeys saw a series of stimulus videos. (**B**) The combinations of cues and stimuli result in a total of 64 types of stimulus videos. Even when the same combination of stimulus video was selected, the initial positions and movement of the stimulus dots varied across trials. (**C**) The iEEG data was chosen from 140 recording sites randomly per trial. (**D**) Activity data was standardized using a 100-ms time window. The decoding model consists of two components: the front-end is a spike decoder using two-layer dense neural networks and the back-end is a video encoder-decoder (i.e., 3D U-Net). The output is a reconstructed video composed of images over time. (**E**) Pearson correlation of pixels between the reconstructed cue and the ground truth. Each bar represents the average correlation over 16 stimuli for the same cue. (**F**) Correlation (Pearson correlation and cosine similarity) of RGB value between the reconstructed stimuli and the ground truth. (**G**) Comparison of the movement direction between the reconstructed video and the ground truth. The angle difference across frames is presented, with the dashed line representing the ground truth angle and the dots indicating the estimated angle from reconstructed videos.

Based on the aforementioned hypothesis, brain data from trained monkeys exhibits similar spike patterns when exposed to familiar stimulus videos, such as stimuli with the same color and direction though moving at different speeds. Given the recognized plausibility of vision-like convolutional neural networks (CNNs) [22, 23, 24, 25, 26, 27, 28], we employed a decoding model that integrates a spike decoder and 3D U-Net architecture to reconstruct dynamic videos featuring moving colored stimulus dots. To foreshadow our results, we discerned comprehensive reconstruction information across six brain regions. The decoding model exhibited strong performance in reconstructing visual videos, particularly within regions associated with vision, while regions unrelated to vision demonstrated limited reconstruction capability. Furthermore, we demonstrated a reciprocal relationship between brain encoding and decoding, as evidenced by the ability of the inverse decoding model to predict neuronal activity from visual input, yielding competitive scores. Additionally, we illustrated that vision-like 3D CNNs would learn patches reflecting visual distinctions in the decoding process [29].

## Results

### Decoding visual stimulus videos from iEEG data

To ensure the impartiality of recording sites, we randomly selected neuronal activity from 140 sites spanning six brain regions for each trial (PFC: 30 sites, FEF: 30 sites, LIP: 30 sites, IT: 10 sites, V4: 20 sites, MT: 20 sites, see Methods). The iEEG data was recorded in the form of discrete spiking times (Fig. 1C). Subsequently, we applied a 100-ms time window and converted the spike times into a standardized activity sequence, serving as the input for our decoding model (Fig. 1D). The hypothesis above suggests that the sequence of neuronal activity harbors concealed patterns representing different types of videos. To validate this hypothesis, we trained the decoding model consisting of a spike decoder as the front-end and a video encoder-decoder component as the back-end. The objective was for the model to generate a corresponding stimulus-type video as output.

Figure 1 shows the model performance in decoding dynamic videos from monkey iEEG data. The video sequence was segmented to evaluate the cue and stimuli stages separately. During the cue stage (from 0.5 to 1.5 s), we calculated the Pearson correlation *C* of the pixels between reconstructed cues and ground truth, regardless of stimuli type (for cross shape: *C* = 0.9986 *±* 0.0001, quatrefoil shape: *C* = 0.9978 *±* 0.0003, circle shape: *C* = 0.9988 *±* 0.0001, and triangle shape: *C* = 0.9977 *±* 0.0002). For the stimuli stage (from 1.5 to 4.5 s), we extracted the RGB color values of the stimulus dots from predicted stimulus videos and ground truth (Pearson correlation: *C* = 0.9959 *±* 0.0023, cosine similarity: *C* = 0.9987 *±* 0.0013). The motion experiment was configured with angles θ = 90^0^, 30^0^, *-*30^0^, *-*90^0^. The predicted motion of stimulus dots across frames was computed using the optical flow method (see Methods), resulting in estimates of θ = 89^0^ *±* 2^0^, 31^0^ *±* 3^0^, *-*29^0^ *±* 2^0^, *-*86^0^ *±* 2^0^, respectively. The iEEG data exhibits high spatio-temporal characteristics and performs well across frequency sampling rates (see Supplemental Files).

### Region-level function convergence between machine and brain

When the machine encounters the same visual stimuli as the brain, functional convergence between them is anticipated. To explore this alignment, presumed to be mediated by representations encoded in neuronal activity, we evaluated the influence of brain areas by applying spike masking (i.e., rest neuronal activity). In this approach, we selectively masked neuronal activity based on its spatial location (region or lobe) and input masked neuronal activity into the trained model to observe the predicted videos (Fig. 2A). To assess the quality of the reconstructed videos, we calculated the pixel differences (e.g., RGB values or movement angles across frames) between scenarios with and without spike masks (Figs. 2B-G). A larger difference value indicates a more pronounced impact of masking the specific area on the quality of reconstruction.

**Fig. 2.**
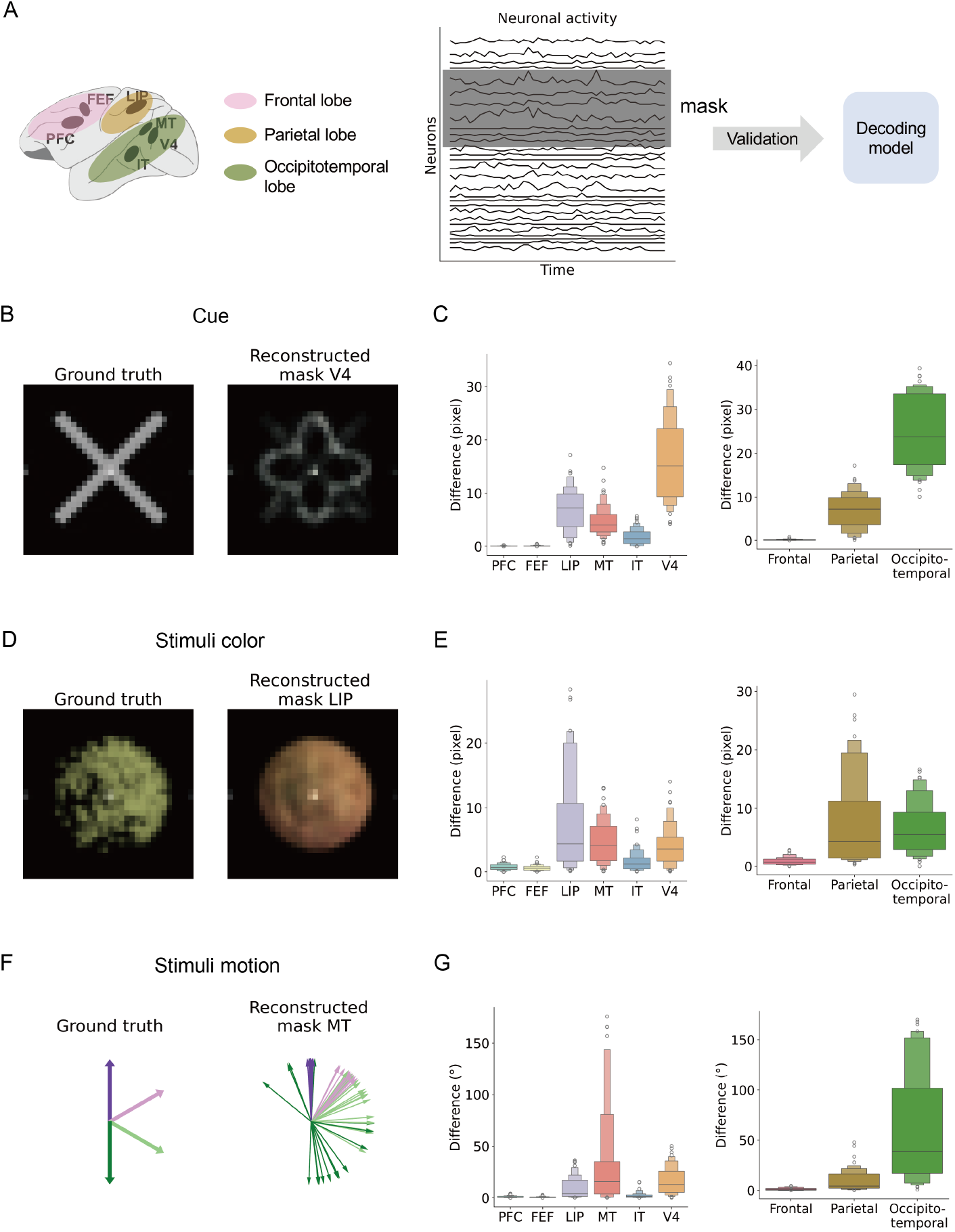
Region-level functional convergence revealed by masked neuronal activity. (**A**) Reconstructing stimulus videos using masked neuronal activity, where the activity in six brain regions (FEF, PFC, LIP, IT, MT and V4) or three lobes (frontal, parietal, occipitotemporal) was individually masked. All the neuronal activity data was used for training and only the unmasked data was used for reconstruction. (**B**,**D**,**F**) Example comparison between the reconstructed and the ground truth highlights the impact of individually masking region V4, LIP and MT on the decoding model’s accuracy regarding shape, color and motion direction respectively. (**C**,**E**,**G**) The difference regarding shape, color and motion direction when individually masking six brain regions and three lobes. Each box represents the average difference over 64 types of videos. The upper and lower boundaries of the box represent the upper and lower quartiles, while the horizontal line inside the box represents the median. Outliers are identified with circles. A larger value indicates that masking the specific area has a greater impact on the reconstruction quality.

We observed a significant influence of vision-related brain areas on the reconstructed visual stimulus videos. Specifically, V4 exhibited a notable impact on the reconstructed shapes of cues, followed by LIP, MT, and IT (mean *±* s.d.: V4: 16.42 *±* 7.83, LIP: 7.11 *±* 4.01, MT: 4.76 *±* 2.87, IT: 1.81 *±* 1.52).

A two-sided *t*-test revealed significant differences in the distributions of pixel (V4: *t*(126) = *-*16.34, *p <* 0.0001, LIP: *t*(126) = *-*12.85, *p <* 0.0001, MT: *t*(126) = *-*11.99, *p <* 0.0001, IT: *t*(126) = *-*7.42, *p <* 0.0001). PFC and FEF did not significantly affect the reconstruction of cues (see Supplemental Table S1). We identified LIP, MT, V4, and IT as influential in shaping the reconstructed color quality of stimulus dots (mean *±* s.d. (RGB value) for LIP: 8.57 *±* 8.52, MT: 2.98 *±* 2.33, V4: 3.55 *±* 2.65, and IT: 1.07 *±* 0.93). LIP was the significant influencer towards reconstructed color (*t*(126) = *-*3.07, *p* = 0.0026, see Supplemental Table S2). Remarkably, MT emerged as a highly influential region for motion direction, with contributions also observed from V4 and LIP (mean *±* s.d. (angle degree) for MT: 30.75 *±* 40.48, *p* = 0.0014, V4: 15.88 *±* 12.61, *p<* 0.0001, and LIP: 9.18 *±* 9.06, *p<* 0.0001, two-sided *t*-test statistic in Supplemental Table S3). These findings underscore the intricate interplay of cortical regions – LIP, V4, MT, and IT – in shaping visual perceptions, ranging from visual categorization to motion processing and recognition of complex visual stimuli.

### CNN representation captures the implicit attention of stimuli

The intricate nature of recorded neuronal activity poses significant challenges in effectively discerning visual videos when subjected to dimensionality reduction on a two-dimensional plane. As shown in Fig. 3A and B, simply feeding the neuronal activity directly into the dimension-reduction method t-SNE (tdistributed stochastic neighbor embedding [30]) fails to adequately discriminate different types of cues and stimuli. However, when leveraging the representation captured by a 3D U-Net (i.e., blue patches in Fig. 1D), the convolutional neural network (CNN) demonstrates remarkable efficacy in accentuating four cues and segregating 16 stimuli at each cue’s location. This finding is substantiated by analogous results obtained with another feature decomposition method UMAP (uniform manifold approximation and projection [31]) in Supplemental Figure S1. These results suggest that CNN harbors the capability to capture implicit attentional representations crucial for discriminating stimuli across diverse cue contexts. CNN exhibits proficiency in reconstructing distinct stimulus videos, underscoring their potential utility in decoding complex neuronal dynamics.

**Fig. 3.**
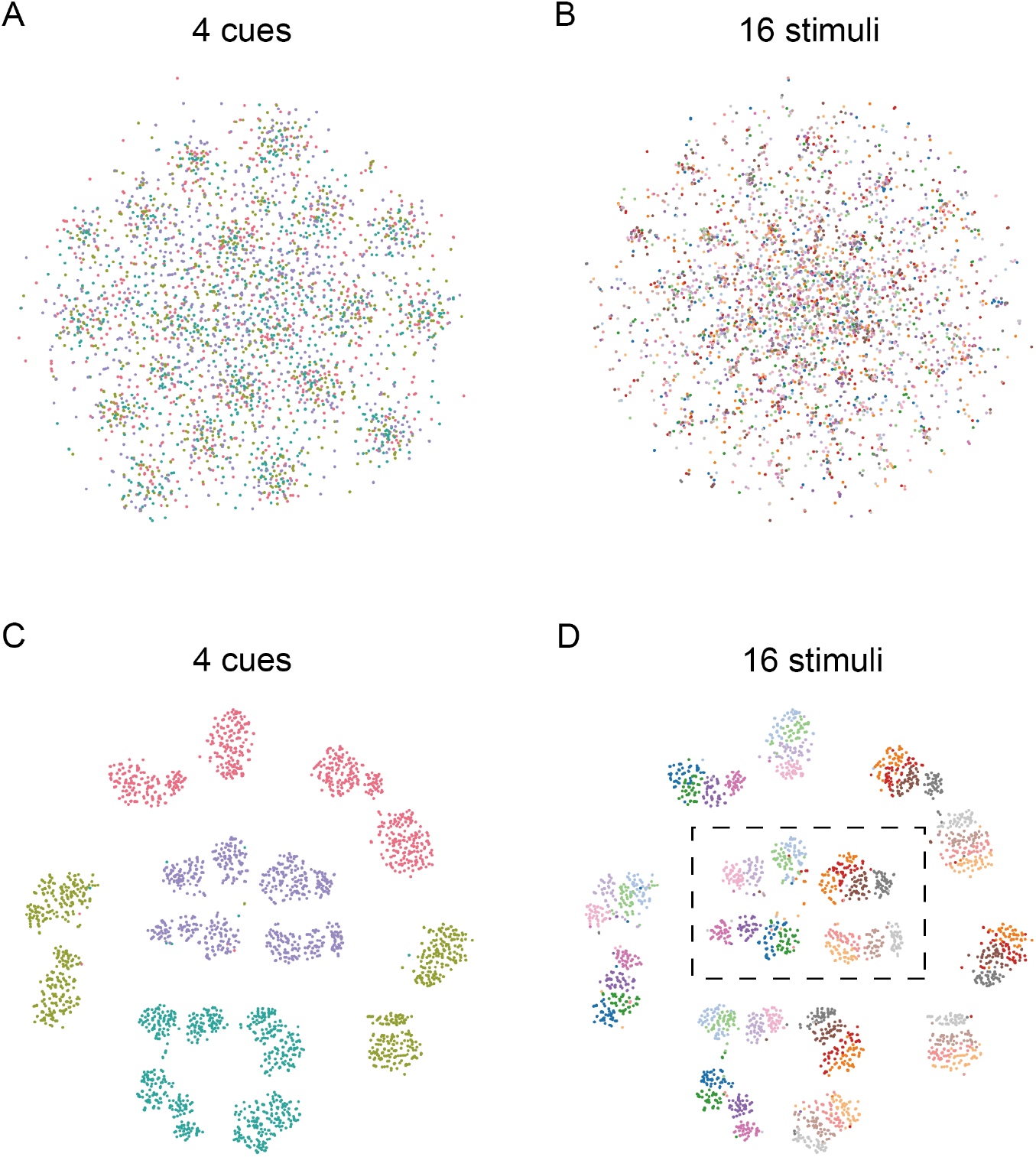
Low-dimensional decomposition of iEEG data. (**A, B**) The t-SNE visualization on a twodimensional plane using direct input of recorded iEEG data. Each category is marked by a unique color. For instance, the four types of cues are represented by four distinct colors (A), while the 16 stimuli are marked by 16 colors (B). (**C, D**) The t-SNE visualization by inputting the model representation (i.e., patches) extracted from the encoder. These two panels distinctly illustrate four colors representing the cues (C), with each cue corresponding to 16 stimulus categories, as highlighted by the dashed black box in (D).

### Decoding dynamic images from individual brain areas

The results of visual decoding from the neuronal activity in multiple brain regions demonstrate impressive performance. However, the question remains: is there redundancy in the brain’s encoding of visual stimuli? In simpler terms, can the stimulus video be decoded from individual areas rather than requiring input from multiple areas collectively?

Therefore, we further evaluate the decoding performance of reconstructed visual videos from individual brain regions or lobes, by training the decoding model with neuronal activity data from limited areas. As depicted in Fig. 4, the stimulus videos cannot be well-reconstructed if only the neuronal activity in the PFC or FEF is utilized. In contrast, each of the other four regions can be leveraged to reconstruct videos that exhibit a high correlation with the ground truth (mean: LIP: *C* = 0.9986, *p<* 0.0001, MT: *C* = 0.9985, *p<* 0.0001, IT: *C* = 0.9977, *p<* 0.0001, V4: *C* = 0.9986, *p<* 0.0001, see Supplemental Table S7). While PFC and FEF appear to recognize the color scheme, they lack the ability to generate refined color details (Fig. 4B). Neural activity in individual LIP, MT, IT, or V4 can reconstruct detailed colors (mean: LIP: *C* = 0.9967, *p <* 0.0001, MT: *C* = 0.9997, *p <* 0.0001, IT: *C* = 0.9930, *p <* 0.0001, V4: *C* = 0.9982, *p<* 0.0001, see Supplemental Table S8). Regarding motion, as shown in Fig. 4C, PFC and FEF appear capable of distinguishing upward and downward directions but are less effective in predicting four directions (i.e., with a high error bar, PFC: s.d.= 25.32^0^, FEF: s.d.= 34.88^0^, see Supplemental Table S9). We also performed 100 independent runs of visual reconstruction from random noise (see Methods), and compared them with that reconstructed from each brain region. As shown in Supplemental Tables S7-9, the results indicate that PFC and FEF cannot fully reconstruct but still contain partial information on the cues or stimuli.

**Fig. 4.**
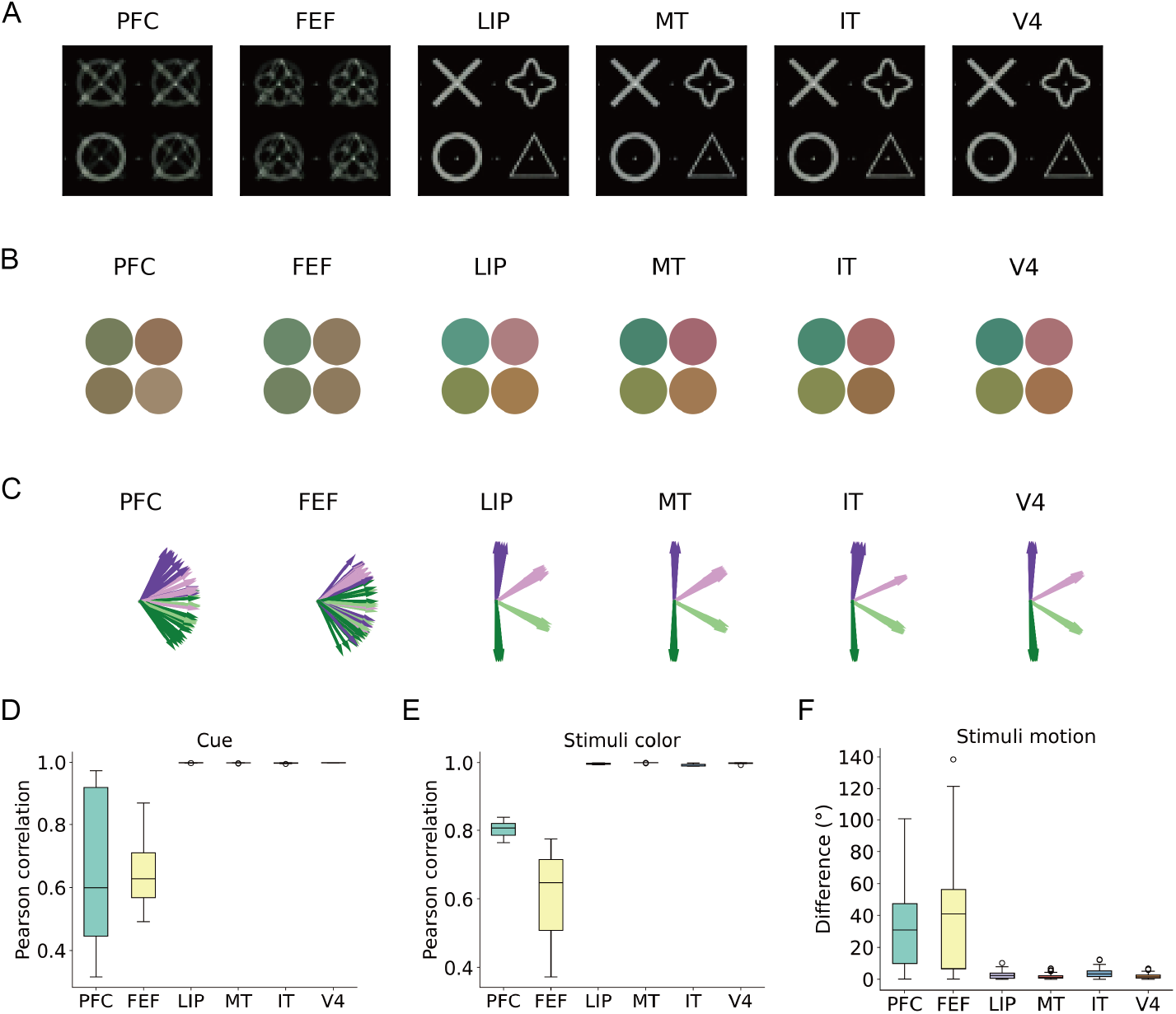
Decoding from brain activity in individual region. (**A**) Reconstructed cues based on neuronal activity in six brain regions individually. A high-quality reconstruction would showcase distinct cues, such as those from individual region LIP, MT, IT, or V4. (**B**) The color of stimulus dots extracted from the reconstructed stimuli. The left side features a greenish color scheme, while the right side reddish. The ground truth comprises four distinct colors, as observed in the reconstruction from LIP, MT, IT, or V4. (**C**) The motion of stimulus dots estimated from the stimulus video across frames. A high-quality reconstruction would reveal four distinct directions, as shown from LIP, MT, IT, and V4. (**D**) Pearson correlation between the ground truth and the reconstructed cues (A). Each box represents the average correlation over 64 types of videos. The upper and lower boundaries of the box denote the quartiles, and the horizontal line inside the box denotes the median. Outliers are marked by circles. A higher correlation value indicates a stronger resemblance to the ground truth. (**E**) Similar evaluation for RGB value based on reconstructed stimuli (B). Higher correlation indicates better performance. (**F**) The angle difference between the ground truth and the reconstructed stimuli motion (C). Smaller difference indicates superior performance.

### Encoding and decoding are reciprocal processes

The CNN model is able to decode the dynamic images from neuronal activity as demonstrated above. We are then interested in the question of whether encoding and decoding are two reciprocal processes, i.e., whether an inverse version of the model can encode the visual stimuli. While previous studies employed distinct models for decoding and encoding brain activity respectively [9, 32], here we simply inverted the CNN model to generate neuronal activity for each cue or stimulus and compared the predicted activity to the ground truth (Fig. 5A).

**Fig. 5.**
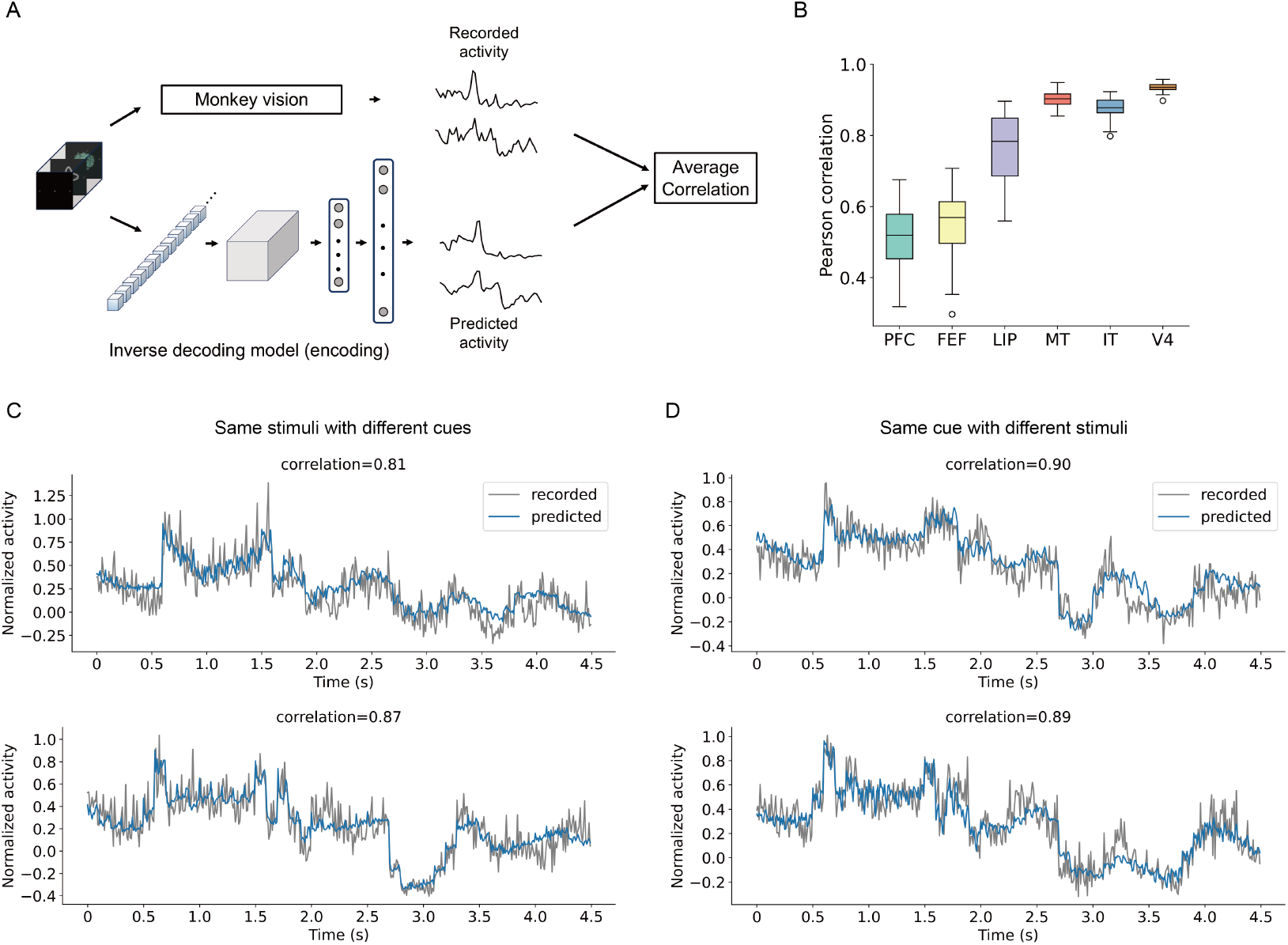
Performance of the inverse decoding model. (**A**) Recorded activity refers to the neuronal recordings when monkeys watched videos, and predicted activity is the output of the inverse decoding model. The similarity between the recorded and the predicted activities is quantified by average Pearson correlation. (**B**) Average Pearson correlation for each of the six brain regions. A higher correlation indicates a closer resemblance between predicted and recorded activities in the specific region. Black line in box denotes median and circles denote outliers. (**C**) Examples of recorded (grey) and predicted (blue) activities in region IT. The monkey was shown the same type of stimulus combination but different cues. (**D**) Examples of recorded (grey) and predicted (blue) activities in region IT. The monkey was shown different stimulus combination but the same type of cue.

The results in Fig. 5B show that the correlation between the predicted and the true neuronal activity is high for vision-related regions LIP, MT, IT, and V4 (mean: LIP: *C* = 0.7671, MT: *C* = 0.9016, IT: *C* = 0.8772 and V4: *C* = 0.9352, see Supplemental Table S10). Such high performance was achieved by the inverse decoding model, suggesting that decoding and encoding are indeed reciprocal. In contrast, the correlation is relatively lower for PFC and FEF (mean: PFC: *C* = 0.5146 and FEF: *C* = 0.5476) which are less relevant to visual information recessing. Moreover, we conducted 100 independent tests by using random noise as input, finding that, even in vision-unrelated areas such as PFC and FEF, the correlation is significantly higher than random prediction (PFC: *t*(162) = 7.624,*p <* 0.0001 and FEF: *t*(162) = 2.078,*p* = 0.0391, see Supplemental Table S10). This result suggests that PFC and FEF may encode partial information of vision, but such encoding appears to be limited, possibly due to functional constraints. Fig. 5C-D displayed typical examples of the neuronal activity recorded and predicted under different stimuli and cues.

## Discussion

This study demonstrated the sophisticated ability of an artificial neural network (ANN)-based machine to decode stimulus videos from iEEE recordings of monkey brains. The machine accurately reconstructed fine features, including shape, color, and motion direction, in the videos. This ability is primarily attributed to the machine’s capacity to compress the original recorded activity into small-scale patches, effectively delineating the hidden connections between brain activities and external stimuli.

We investigated the association between decoding ability and specific brain regions. Instead of using activity data from the entire brain, we focused on data from individual regions for decoding. The results revealed that stimulus videos were best reconstructed when utilizing activity data from vision-related regions, including LIP, MT, IT, and V4. In contrast, decoding quality was lower when using only activity data from PFC and FEF regions, which are presumed to be less involved in visual functions. This discovery suggests that task-unrelated regions may encode partial information about external stimuli, but their activities alone are insufficient for effective decoding. Such strategic utilization of regionspecific information has the potential to further optimize the decoder and enhance the proficiency of brain-machine interfaces.

We inverted the machine architecture to predict the brain activity given various stimulus videos, finding that decoding and encoding are part of a mutual process. The observed impact of brain regions remains consistent in the inverse process. Although we could encode visual stimuli in multiple brain regions, the performance was limited and less effective when encoding dynamic images in vision-unrelated brain regions. These insights deepen our understanding of the intricate relationship between artificial intelligence and cognitive processes.

Our study has several limitations. Firstly, our focus was solely on the correspondence between visual stimuli and neuronal activity, without considering decision-making based on monkey eye movements. This may explain the underutilization of PFC and FEF brain regions. Previous work [17] has shown that when the machine encounters cognitive decision-making pressure and needs to respond with an eye saccade, a corresponding relationship with the PFC and FEF brain regions would emerge. Therefore, we argue that under varying task pressures, the machine acquires distinct regional functions, particularly exhibiting a high correlation with brain areas related to vision when confronted with visual pressure.

Secondly, the brain’s ability to evoke similar neural patterns in familiar environments may reflect its remarkable plasticity and adaptability. In familiar settings, monkeys develop routines that trigger specific neural patterns. An ANN-based machine effectively decoded these patterns, aligning with the intended regional function. However, when faced with environmental changes, the brain likely adjusts its activity to suit new circumstances. Particularly in unfamiliar or ambiguous situations, deciphering neural activity becomes challenging.

## Methods

### Monkey experiment

Two rhesus monkeys, one female and one male, were trained on a flexible visuomotor decision-making task over the years. The visuomotor decision-making task rewarded animals for providing correct responses based on the presented stimulus videos. We analyzed the neuronal activity recorded from these rhesus monkeys exposed to a series of stimulus videos after training. The stimulus videos presented to monkeys during both training and recording sessions consisted of three segments: a fixation period (0 s to 0.5 s), a cue period (0.5 s to 1.5 s), and a stimuli presentation period (1.5 s to 4.5 s). Depending on the task cued at the beginning of each trial, the monkeys categorized either the color (red vs. green) or motion direction (up vs. down) of the stimulus and reported their perception with a left or right saccade. In this study, our focus was on analyzing the visual stimuli and corresponding neuronal activity, independent of saccade categorization. A new stimulus was generated for each recording trial. The stimulus consisted of randomly positioned colored dots moving in a coherent direction. There were four possible colors and four possible motion directions (see Fig. 1). Although the stimuli set consists of 16 color-motion items, each stimulus was randomized due to the initial random positions. For specific details regarding the stimulus images, please refer to Supplementary Information.

### Invasive EEG data

Each monkey was implanted with a titanium head bolt to immobilize the head. Recordings were performed through three recording chambers that were stereotactically placed over the frontal (including FEF and PFC), parietal (including LIP), and occipitotemporal (including MT, V4, and IT) cortex. Multi-unit spiking activity (MUA) was recorded from 2694 recording sites (1753 and 941 for two monkeys) in 48 sessions (31 and 17 for the two monkeys) and each recording session consists of hundreds of trials. The stimuli in each trial were randomized. Electrodes were advanced simultaneously in pairs or triplets, with penetrations typically angled relative to the cortical surface. There was no specific targeting of layers or fine-tuning of particular cells.

### Data processing

We used the iEEG data for two monkeys after training as monkeys often took years to train without training data (i.e., spike trains) available. In the original experiment [18], neuronal activity was recorded at least from three regions simultaneously for each session. To ensure the impartiality of recording sites, we considered recording sessions with all six investigated regions. Considering the area and number of electrode placements across the six brain regions, we randomly selected a total of 140 sites for each trial (PFC: 30 sites, FEF: 30 sites, LIP: 30 sites, IT: 10 sites, V4: 20 sites, MT: 20 sites). We then considered the activity sequence from 140 different recording sites (with less than 10% overlap) as a sample trial and extracted 512 sample trials that met the requirements for each cue and stimulus combination. Neuronal activity was obtained using rate coding (i.e., spike count) with a 100-ms time window. We adopted a training/test/evaluation ratio of 7:2:1. Prior to training the model with the iEEG dataset, we performed standardization to preserve its distribution.

### Video processing

Each trial was initiated when the monkey fixated on a point at the center of the screen and required fixation within 1.2^0^ of visual angle from the fixation point, which was maintained throughout. Following a fixation period (500 ms), the monkey was presented with a visual task cue lasting 1000 ms. The cues, depicted in Fig. 1B, consisted of one of four distinct gray shapes, each approximately 1.5^0^ in visual angle diameter, and were centrally presented on the fixation spot. After the cue period, the cue disappeared, and stimulus dots appeared at the center of the fixation spot. These were images of random colored dots, all moving synchronously in a specific direction. The random dots appeared within a circle centered at the fixation point. All stimuli were 100% coherent, combining one of four possible colors and one of four possible motions. Due to the randomness of position and velocity, the stimuli exhibited fluctuation in color direction within a circular region when averaged or integrated. Hence, we hypothesized that the machine is capable of learning averaged color and motion across trials.

### Decoding model

To reconstruct chromatic and continuous vision from iEEG data, we employed a decoding model, consisting of a spike decoder denoted as **f**, and a video encoder-decoder, as **g** and **h**, respectively. These components work cohesively to decode spikes into hidden features and then generate the corresponding video output.

The spike decoder utilizes a two-layer fully connected neural network to transform iEEG data *x* ∈ ℝ^N×T^ to images using the Unflatten operation,

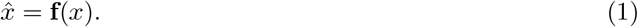

where *N* = 140 sites and *T* = 45 time points corresponding to a 100-ms bin within 4500 ms. Batch normalization and ReLU activation between layers were implemented to ensure effective training and introduce non-linearity, aiding the transformation from iEEG data to images.

CNNs are inspired by human visual neurons [33]. The video encoder-decoder is a 3D U-Net, including repetitions of many convolution layers. The video encoder is to extract hidden features (i.e., patches) from the spike-decoder, while the video decoder aims to reconstruct the video from these patches. Specially, this video encoder contains four repetitive convolutional layers with kernel size of 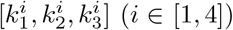, followed by four batch normalizations and ReLU activations, and max poolings with kernel size of 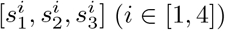, and then generates patches *c*,

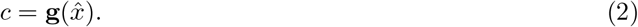

The convolutional kernel size is 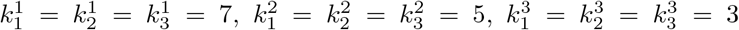 and 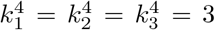, and the stride is of 1. The max pooling with kernel size of 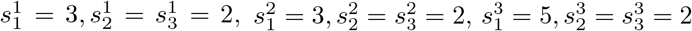, and 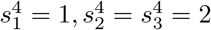, respectively.

The video decoder, which is a reflection of the encoder, encompasses four convolutional layers, batch normalizations, ReLU activations, and up-samplings, yielding an output *v* ∈ ℝ^channels,T,height,width^,

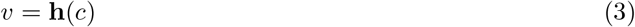

where channels, height, and width are 3, 32, and 32, respectively. Here we used an image of 64 × 64 pixels but cropped it into 32 ×32 due to the black background surrounding it (see Supplemental Information). Finally, we employed the hyperbolic tan function to regulate the output 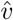 within the range of [0, 1],

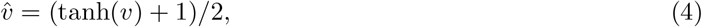

and

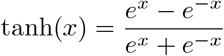

thus elegantly transforming iEEG brain data to RGB-based videos.

### Loss function

The loss function employed in our model comprises two components: the Structural Similarity Index Measure (SSIM) that encourages to preserve structural information, and the Mean Squared Error (MSE) that minimizes the average squared differences between corresponding pixels. This hybrid approach is crafted to encompass both the perceptual similarity between images and the pixel-wise intensity differences. Formally, the loss function is as follows:

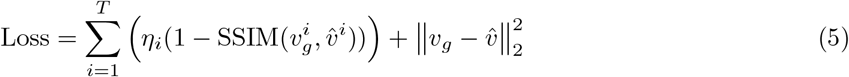

where 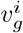 denotes the *i*-th image in the ground truth video, 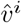 indicates the *i*-th image from reconstructed video, and η is the weight assigned to different images. We chose η = 1, 2, 5 for fixation, cue, and stimulus images, respectively. For more detailed information about the loss function, please see Supplemental Information.

In this study, we selected a batch size of 64 and employed the Adamax optimizer with an initial learning rate of 0.01 because of its robustness in handling sparse gradients. Additionally, to enhance the training process and achieve better performance, we implemented a StepLR scheduler [34], which reduced the learning rate by a factor of 0.85 every 3 epochs.

### Evaluating reconstructed videos

#### Cue images

The cue images could be one of four shapes: cross, quatrefoil, circle, or triangle. These cues appeared between 500 ms and 1500 ms after the fixation started at 0 ms. Since the cue in the stimulus video remains static, we directly assessed the Pearson correlation of pixels between the reconstructed and original cue images.

### RGB color of stimulus images

As the stimulus points move in a specific direction over time, color variations occur at consistent pixel positions across frames. Considering the randomized initial positions on each trial, we computed aggregated type-based stimulus images across trials, resulting in stimulus dots arranged within a circle. The RGB values may slightly deviate from the assigned colors due to the black background. Specifically, we selected four symmetric points located at 2/4 and 3/4 positions of the aggregated stimulus images to evaluate the correlation between predicted stimulus dots and the ground truth based on their respective RGB values.

### Movement across stimulus images

To assess the movement of stimulus dots over time, we conducted optical flow calculations on a sequence of images. This entailed iterating through frames, converting each frame image to grayscale, and computing optical flow using the Farneback method. The averaged motion vector was derived from the optical flow matrix over consecutive frames with a step of 2. This algorithm is implemented in OpenCV as `calcOpticalFlowFarneback` [35].

### Visualization of high-dimensional data

To illustrate the complexity of recorded neuronal activity, we employed two unsupervised dimensionality reduction and manifold learning techniques: t-Distributed Stochastic Neighbor Embedding (t-SNE) [30] and Uniform Manifold Approximation and Projection (UMAP) [31] (see Supplemental Information). These methods facilitate the decomposition of high-dimensional data without the need for labeled information. The visualized results were presented by reducing the high-dimensional data to a two-dimensional plane.

### Statistical analyses

We evaluated the statistical significance of the comparison results. To discern the function of each individual region learned by the model, we conducted a two-sided *t*-test comparing the predicted pixels reconstructed using rest-masked iEEG data to their corresponding predicted pixels without masks. Additionally, to assess the statistical significance of predicted videos and neuronal activity in each individual brain area, we conducted 100 independent tests using random Gaussian input *N* (0, 1). Specifically, we tested random activity as input in the decoding process, while random pixels were used as input in the encoding process. The p-values were computed as the fraction of null sequences with predicted pixels/activities differing from the output. All p-values were indicated as follows: ^*^*p <* 0.05, ^* *^*p <* 0.01,^* * *^*p<* 0.001, ^* * * *^*p<* 0.0001.

## Acknowledgements

This work was supported by the National Natural Science Foundation of China (Grants No. T2225022, No. 12161141016, and No. 62088101), Shuguang Program of Shanghai Education Development Foundation and Shanghai Municipal Education Commission (Grant No. 22SG21), Shanghai Municipal Science and Technology Major Project (Grant No. 2021SHZDZX0100), Shanghai Municipal Commission of Science and Technology Project (Grant No. 19511132101), and the Fundamental Research Funds for the Central Universities. We modified the photo of a monkey brain depicted in Fig. 1 and Fig. 2, with credit to Macauley Smith Breault at https://doi.org/10.5281/zenodo.3910249.

## Data and Code availability

The codes used in this work are available at https://github.com/ganglab/NeuralDecodingVideo. All behavioral and electrophysiological data are archived at the Centre for Integrative Neuroscience, University of Tübingen, Germany.

## Supplementary Information for

## I Materials and Methods

### Monkey experiment

Two rhesus monkeys, a female named Paula, and a male named Rex, underwent training for a categorization task. The animals had to respond with one direct saccade that categorized either the color (red vs. green) or motion direction (up vs. down) of the stimulus based on the task cue. For correct responses, the animals were rewarded with apple juice. After years of training, neuronal activity was recorded in six brain regions. In each recording session, all pairs of regions were recorded simultaneously at least once. Each session comprised multiple trials, where each trial consisted of a sequence of animated images, including a fixation (lasting 0.5 s), a cue (lasting 1 s), and stimuli (lasting 3 s). The cue could be one of four types: cross, quatrefoil, circle, or triangle shapes. After the cue period, the cue was switched off and stimuli appeared at the center of the fixation spot. Neuronal activity was recorded across brain areas simultaneously throughout the videos. The videos featured a static fixation point and cue images, while the stimulus images were dynamic, consisting of 400 moving dots.

### Stimulus image

Stimulus images were 64 *×* 64 RGB pixels. In the fixation image, the distance between the fixation point and either of the grey dots is equal to 1*/*4 of the horizontal length of the image. This ratio is kept constant for the rest of the image types. Monochrome cue images were imported, the cue color changed to grey, the background to black, and three dots (also present in the fixation point image) added. The stimuli are dynamic random dot patterns with 100% motion coherence, centered on the fixation spot. The stimuli have a diameter of 3.2, a dot diameter of 0.08, 400 dots, and two dot speeds (1.67 /s or 10 /s) for half of the recording trials. We considered four possible colors and four possible directions for the stimulus dots. The direction labels are −90°, −30°, 30°, 90°. All colors are defined in the CIE Lab* space with identical luminance and saturation. In total, there were 16 possible color-motion combinations presented in Fig. 1. Due to the redundant black background in the video surroundings, we cropped the 64 *×* 64 pixels to 32 *×* 32 pixels by removing the surrounding black background.

### Decoding model

To train our decoding model, we replicated fixation and cue images from the original experiment and generated stimulus images with the above-mentioned stimulus patterns. As the fixation point and cue images were static, we repeated them to simulate the movies presented to the monkeys. Considering an *f* -ms window for processing iEEG data (equivalent to a frequency of 1*/f* frames per second), each trial comprises 4.5 *×* 1*/f* images, including 0.5 *×* 1*/f* fixation point images, 1.0 *×* 1*/f* cue images, and 3.0 *×* 1*/f* stimulus images. In this study, the time bin/frequency parameter *f* was selected to ensure an integer result for the specified number of images mentioned above.

### Loss function

The loss function employed in our model comprises two components: the Structural Similarity Index Measure (SSIM) and the Mean Squared Error (MSE). For any images *x* and *y*, the objective function is to minimize

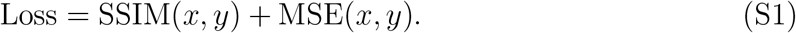

The SSIM component encourages the model to preserve structural information. For images *x* and *y*, SSIM is calculated using the following formula:

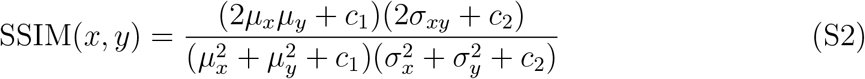

where μ_x_ and μ_y_ represent mean intensities of images *x* and *y*, 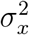 and 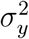 are the variances of *x* and *y*, σ_xy_ denotes the covariance between *x* and *y*, and *c*_1_, *c*_2_ are the constants to avoid instability near zero. The SSIM ranges from -1 to 1, where 1 indicates perfect similarity between images. Therefore, we utilized *η* · (1 − SSIM(*x, y*)) in our study, where *η* denotes the importance weight assigned to different images. Specifically, we set *η* to 1, 2, and 5 for fixation, cue, and stimulus images, respectively.

The MSE term ensures that the model minimizes the mean squared differences between corresponding pixels,

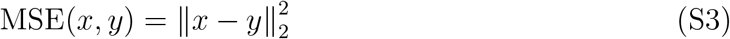

thereby increasing the overall color accuracy of the reconstructed images.

### Estimating optical flow using Farneback method

To compute the optical flow, we employed the Farneback method. Initially, we iterated through each frame of the video, converting it to grayscale using the OpenCV cvtColor function. The optical flow was then computed using the Farneback method, implemented through the `calcOpticalFlowFarneback` function in OpenCV [1],

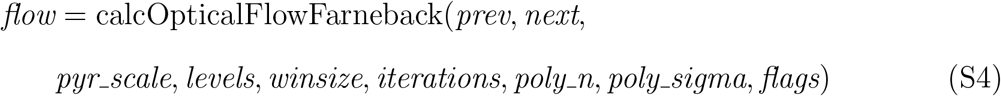

where *prev* and *next* represent the input frames (i.e., the grayscale images), and *flow* denotes the resulting optical flow vector. The algorithm parameters, including *pyr scale, levels, winsize, iterations, poly n, poly sigma, flags*, are set to 0.5, 3, 15, 3, 5, 1.2, and 0, respectively.

Finally, we obtained the averaged motion vector from the optical flow matrix over consecutive frames with a specified step (e.g., a step of 2). Through these steps, we successfully applied the Farneback method to calculate the optical flow of objects within a video.

### Visualization of high-dimensional data

We utilized two unsupervised dimensionality reduction and manifold learning techniques, t-Distributed Stochastic Neighbor Embedding (t-SNE, Fig. 3) and Uniform Manifold Approximation and Projection (UMAP, see Fig. S1).

### T-distributed stochastic neighbor embedding

The visualization t-SNE is implemented with TSNE [2] in scikit-learn toolbox, with a maximum iteration of 10^3^.

### Uniform manifold approximation and projection

The visualization is implemented with umap-learn toolbox [3] for general non-linear dimension reduction, with a minimum distance between data points of 3*×*10^−1^ and constraining the size of the local neighborhood of 5 neighbors.

## II Tables

**Statistics in Fig. 2**

**Table S1:** The difference (mean*±*s.d.) of pixels for cue images reconstructed from data with masked brain regions. A larger value indicates that masking the specific region has a greater impact on cue reconstruction. A two-sided *t*-test was performed to compare the predicted pixels reconstructed with the masked region to their corresponding predicted pixels.

| Masked region (cue) | mean | s.d. | $t$-test |
| --- | --- | --- | --- |
| PFC | 0.0584 | 0.0583 | $t(126)=-0.0171$, $p=0.9864$ |
| FEF | 0.0832 | 0.1032 | $t(126)=0.4191$, $p=0.6758$ |
| LIP | 7.1086 | 4.0081 | $t(126)=-12.85$, $p<0.0001$ |
| MT | 4.7556 | 2.8706 | $t(126)=-11.99$, $p<0.0001$ |
| IT | 1.8100 | 1.5199 | $t(126)=-7.424$, $p<0.0001$ |
| V4 | 16.416 | 7.8273 | $t(126)=-16.34$, $p<0.0001$ |

**Table S2:**
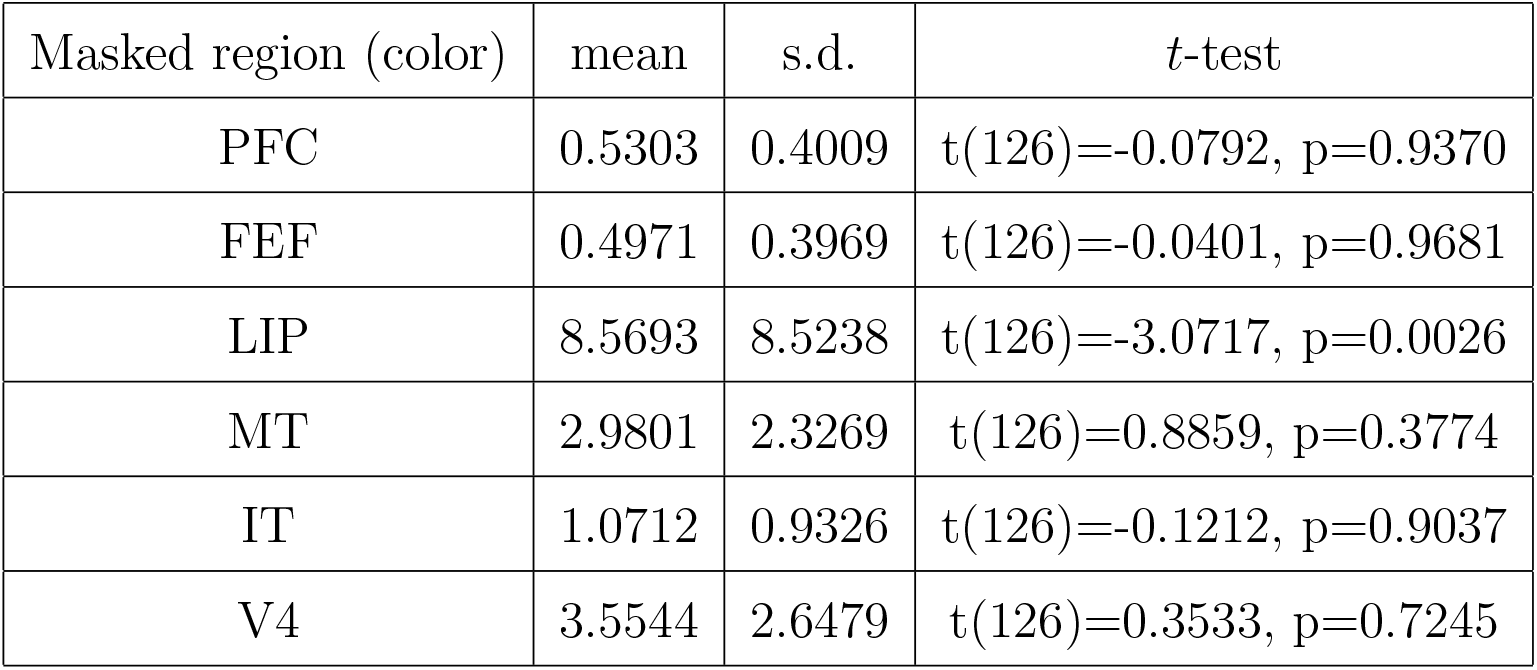
The RGB color difference (mean*±*s.d.) of pixels for stimulus images reconstructed from data with masked brain regions. A larger value indicates that masking the specific region has a greater impact on the reconstruction of stimulus color. A two-sided *t*-test was performed to compare the predicted pixels reconstructed with the masked region to their corresponding predicted pixels.

| Masked region (color) | mean | s.d. | $t$-test |
| --- | --- | --- | --- |
| PFC | 0.5303 | 0.4009 | $t(126)=-0.0792$, $p=0.9370$ |
| FEF | 0.4971 | 0.3969 | $t(126)=-0.0401$, $p=0.9681$ |
| LIP | 8.5693 | 8.5238 | $t(126)=-3.0717$, $p=0.0026$ |
| MT | 2.9801 | 2.3269 | $t(126)=0.8859$, $p=0.3774$ |
| IT | 1.0712 | 0.9326 | $t(126)=-0.1212$, $p=0.9037$ |
| V4 | 3.5544 | 2.6479 | $t(126)=0.3533$, $p=0.7245$ |

**Table S3:** The movement angle difference (mean*±*s.d.) across frames (i.e., stimulus images) reconstructed from data with masked brain regions. A larger value indicates that masking the specific region has a greater impact on the reconstruction of stimulus motion. A two-sided *t*-test was performed to compare the predicted pixels reconstructed with the masked region to their corresponding predicted pixels.

| Masked region (motion) | mean | s.d. | $t$-test |
| --- | --- | --- | --- |
| PFC | 0.9892 | 0.8283 | $t(126)=-0.4301$, $p=0.6678$ |
| FEF | 0.7911 | 0.6103 | $t(126)=0.0993$, $p=0.9210$ |
| LIP | 9.183 | 9.063 | $t(126)=-5.532$, $p<0.0001$ |
| MT | 30.75 | 40.48 | $t(126)=-5.711$, $p=0.0014$ |
| IT | 2.677 | 2.792 | $t(126)=-1.590$, $p=0.1144$ |
| V4 | 15.88 | 12.61 | $t(126)=-8.456$, $p<0.0001$ |

**Table S4:** The difference (mean*±*s.d.) of pixels for cue images reconstructed from data with masked brain lobes. A larger value indicates that masking the specific lobe has a greater impact on cue reconstruction. A two-sided *t*-test was performed to compare the predicted pixels reconstructed with the masked lobe to their corresponding predicted pixels.

| Masked lobe (cue) | mean | s.d. | $t$-test |
| --- | --- | --- | --- |
| Frontal | 0.1337 | 0.1455 | $t(126)=-0.6474$, $p=0.5185$ |
| Parietal | 7.1086 | 4.0081 | $t(126)=-12.8542$, $p<0.0001$ |
| Occipitotemporal | 24.752 | 8.4836 | $t(126)=-22.5927$, $p<0.0001$ |

**Table S5:** The RGB color difference (mean*±*s.d.) of pixels for stimulus images reconstructed from data with masked brain lobes. A larger value indicates that masking the specific lobe has a greater impact on the reconstruction of stimulus color. A two-sided *t*-test was performed to compare the predicted pixels reconstructed with the masked lobe to their corresponding predicted pixels.

| Masked lobe (color) | mean | s.d. | $t$-test |
| --- | --- | --- | --- |
| Frontal | 0.8124 | 0.5377 | $t(126)=-0.1165$, $p=0.9075$ |
| Parietal | 8.5693 | 8.5238 | $t(126)=-3.0717$, $p=0.0026$ |
| Occipitotemporal | 7.1552 | 6.2830 | $t(126)=-1.0150$, $p=0.3120$ |

**Statistics in Fig. 4**

**Table S6:** The movement angle difference (mean*±*s.d.) of stimulus images across frames reconstructed from data with masked brain lobes. A larger value indicates that masking the specific lobe has a greater impact on the reconstruction of stimulus motion. A twosided *t*-test was performed to compare the predicted pixels reconstructed with the masked lobe to their corresponding predicted pixels.

| Masked lobe (motion) | mean | s.d. | $t$-test |
| --- | --- | --- | --- |
| Frontal | 1.396 | 1.065 | $t(126)=-0.5281$, $p=0.5983$ |
| Parietal | 9.425 | 10.43 | $t(126)=-5.311$, $p<0.0001$ |
| Occipitotemporal | 66.91 | 54.60 | $t(126)=-9.492$, $p<0.0001$ |

**Table S7:** Pearson correlation (mean*±*s.d.) of pixels between the ground truth and reconstructed cues from individual brain region data. A lower correlation value implies that information from this particular region is less conducive to achieving good performance. To assess the statistical significance, 100 independent tests were conducted using random Gaussian input, and p-values were computed as the fraction of null sequences with predicted pixels differing from random distributions.

| Region (cue) | mean | s.d. | $t$-test |
| --- | --- | --- | --- |
| PFC | 0.6544 | 0.2273 | $t(162)=6.0369$, $p<0.0001$ |
| FEF | 0.6436 | 0.0927 | $t(162)=9.2237$, $p<0.0001$ |
| LIP | 0.9986 | 0.0004 | $t(162)=585.9$, $p<0.0001$ |
| MT | 0.9985 | 0.0005 | $t(162)=706.1$, $p<0.0001$ |
| IT | 0.9977 | 0.0010 | $t(162)=842.5$, $p<0.0001$ |
| V4 | 0.9986 | 0.0004 | $t(162)=755.8$, $p<0.0001$ |

**Table S8:** Pearson correlation (mean*±*s.d.) of RGB values between the ground truth and reconstructed stimulus images from individual brain region data. A lower correlation value implies that information from this particular region is less conducive to achieving good performance. To assess the statistical significance, 100 independent tests were conducted using random Gaussian input, and p-values were computed as the fraction of null sequences with predicted RGB values differing from random distributions.

| Region (color) | mean | s.d. | *t*-test |
| --- | --- | --- | --- |
| PFC | 0.8008 | 0.0261 | t(162)=51.5067, p<0.0001 |
| FEF | 0.5826 | 0.1352 | t(162)=-21.7066, p<0.0001 |
| LIP | 0.9967 | 0.0023 | t(162)=198.0, p<0.0001 |
| MT | 0.9997 | 0.0004 | t(162)=170.9, p<0.0001 |
| IT | 0.9930 | 0.0029 | t(162)=226.1, p<0.0001 |
| V4 | 0.9982 | 0.0011 | t(162)=119.8, p<0.0001 |

**Table S9:** The movement angle difference (mean*±*s.d.) between the ground truth and reconstructed stimulus videos from individual brain region data. A higher difference value implies that information from this particular region is less conducive to achieving good performance. To assess the statistical significance, 100 independent tests were conducted using random Gaussian input, and p-values were computed as the fraction of null sequences with predicted motion differing from random distributions.

| Region (motion) | mean | s.d. | *t*-test |
| --- | --- | --- | --- |
| PFC | 33.01 | 25.32 | t(162)=-10.61, p<0.0001 |
| FEF | 40.25 | 34.88 | t(162)=-5.552, p<0.0001 |
| LIP | 2.739 | 2.334 | t(162)=-244.6, p<0.0001 |
| MT | 1.882 | 1.518 | t(162)=-134.4, p<0.0001 |
| IT | 4.069 | 2.912 | t(162)=-207.8, p<0.0001 |
| V4 | 2.041 | 1.603 | t(162)=-330.1, p<0.0001 |

**Statistics in Fig. 5**

**Table S10:** Pearson correlation (mean±s.d.) between the recorded activity and the activity predicted from the inverse decoding model. A higher correlation value indicates better performance in predicting activity within that region. To assess the statistical significance, 100 independent tests were conducted using random Gaussian input, and p-values were computed as the fraction of null sequences with predicted activity differing from random distributions.

| Encoding | mean | s.d. | *t*-test |
| --- | --- | --- | --- |
| PFC | 0.5146 | 0.0880 | t(162)=7.624, p<0.0001 |
| FEF | 0.5476 | 0.0929 | t(162)=2.078, p=0.0391 |
| LIP | 0.7671 | 0.0932 | t(162)=15.43, p<0.0001 |
| MT | 0.9016 | 0.0204 | t(162)=14.09, p<0.0001 |
| IT | 0.8772 | 0.0280 | t(162)=47.12, p<0.0001 |
| V4 | 0.9352 | 0.0110 | t(162)=80.28, p<0.0001 |

## III Figures

**Figure S1:**
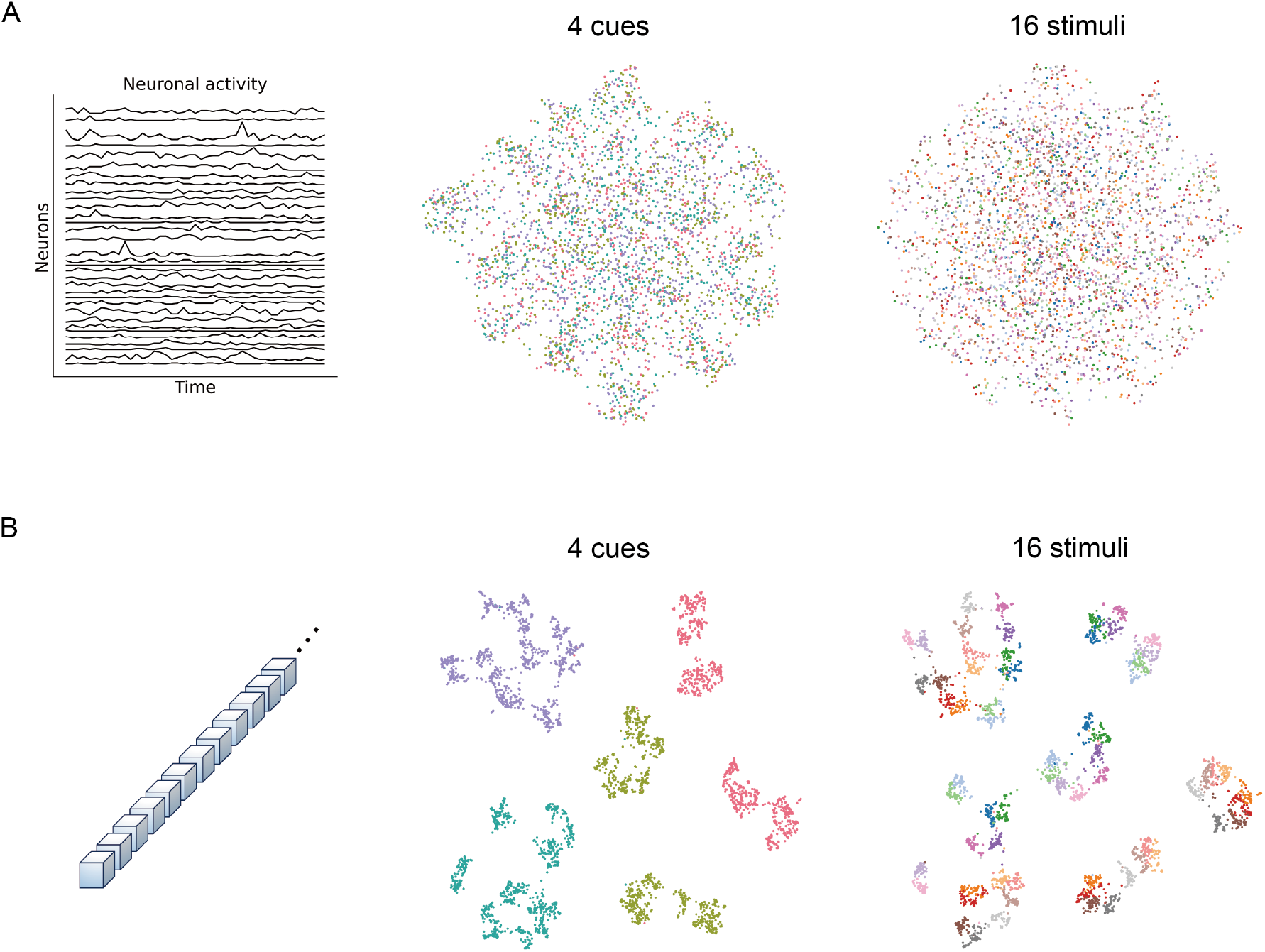
Low-dimensional decomposition of iEEG data. (**A**) The UMAP visualization on a two-dimensional plane using direct input of recorded iEEG data. Each category is marked by a unique color. For instance, the four types of cues are represented by four distinct colors, while the 16 stimuli are marked by 16 colors. (**B**) The UMAP visualization by inputting the model representation (i.e., patches) extracted from the encoder. These panels distinctly illustrate four colors representing the cues, with each cue corresponding to 16 stimulus categories.

**Figure S2:**
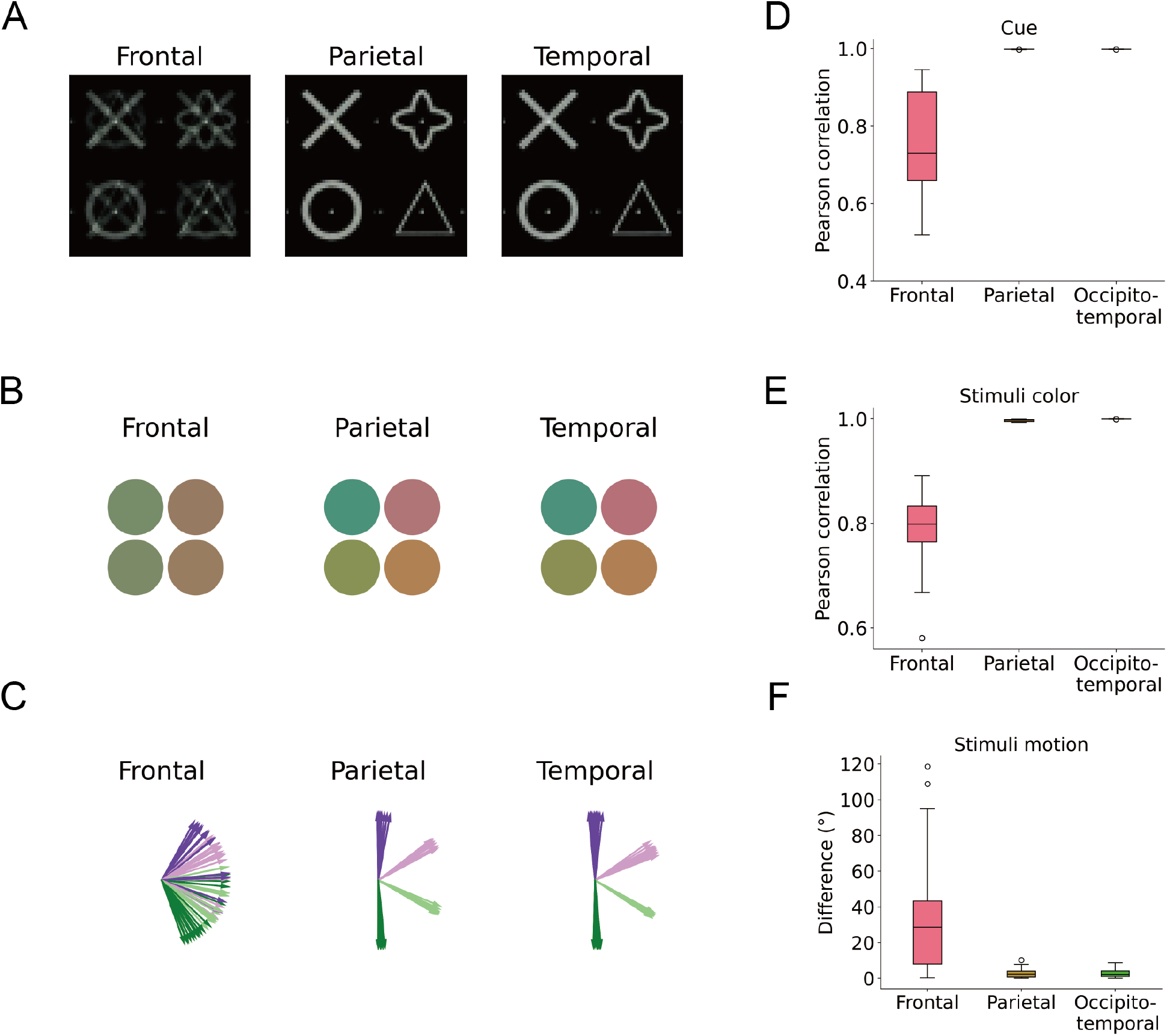
Decoding from brain activity in individual lobe. (**A**) Reconstructed cues based on neuronal activity in three brain lobes individually. A high-quality reconstruction would showcase distinct cues, such as those from individual parietal and occipitotemporal lobe. (**B**) The color of stimulus dots extracted from the reconstructed stimuli. The left side features a greenish color scheme, while the right side features a reddish scheme. The ground truth comprises four distinct colors, as observed in the reconstruction from individual parietal and occipitotemporal lobe. (**C**) The motion of stimulus dots was estimated from the stimulus video across frames. A high-quality reconstruction would reveal four distinct directions, as shown from individual parietal and occipitotemporal lobe. (**D**) Pearson correlation between the ground truth and reconstructed cues (A). Each box represents the average correlation over 64 types of videos. The upper and lower boundaries of the box denote the quartiles, and the horizontal line inside the box denotes the median. Outliers are marked by circles. A higher correlation value indicates a stronger resemblance to the ground truth. (**E**) Similar evaluation for RGB value based on reconstructed stimuli (B). Higher correlation value indicates better performance. (**F**) The angle difference between the ground truth and the reconstructed stimuli motion (C). Smaller difference indicates superior performance.

